# agoTRIBE detects miRNA-target interactions transcriptome-wide in single cells

**DOI:** 10.1101/2022.08.10.503472

**Authors:** Vaishnovi Sekar, Emilio Mármol-Sánchez, Panagiotis Kalogeropoulos, Laura Stanicek, Eduardo A. Sagredo, Evangelos Doukoumopoulos, Franziska Bonath, Inna Biryukova, Marc R. Friedländer

**Author notes:** These authors contributed equally (Emilio Mármol-Sánchez, Panagiotis Kalogeropoulos).

## Abstract

MicroRNAs are gene regulatory molecules that play important roles in numerous biological processes including human health. The function of a given microRNA is defined by its selection of target transcripts, yet current state-of-the-art experimental methods to identify microRNA targets are laborious and require millions of cells. We have overcome these limitations by fusing the microRNA effector protein Argonaute2 to the RNA editing domain of ADAR2, allowing for the first time the detection of microRNA targets transcriptome-wide in single cells. Our agoTRIBE method reports functional microRNA targets which are additionally supported by evolutionary sequence conservation. As a proof-of-principle, we study microRNA interactions in single cells, and find substantial differential targeting across the cell cycle. Lastly, agoTRIBE additionally provides transcriptome-wide measurements of RNA abundance and will allow the deconvolution of microRNA targeting in complex samples such as tissues at the single-cell level.

---

MicroRNAs (miRNAs) are small non-coding RNAs that post-transcriptionally regulate the expression of protein coding genes^1^. Mechanistically, they guide Argonaute effector proteins to mRNA targets, allowing Argonaute and cofactors to inhibit translation and/or promote degradation of the target mRNAs^2-5^. miRNAs are found in virtually all multicellular animals and plants and play important roles in numerous biological processes, including development, formation of cell identity and human diseases such as cancer (reviewed here^1, 6-8^). The human genome harbors hundreds of distinct miRNA genes, each of which can putatively regulate hundreds of target genes. The function of each individual miRNA is defined by its specific target repertoire; thus, to understand the function of a given miRNA, it is necessary to map its targets. The current state-of-the-art method to do so is CLIP-seq, which applies UV light to crosslink the Argonaute protein to its mRNA targets in cells, then isolates the protein using antibodies, and uses next-generation sequencing to profile the bound RNA targets (reviewed here^9, 10^). This method has brought many new insights to the miRNA field, yet it has some inherent limitations. Firstly, since the isolation with antibodies is inefficient, it requires in the order of millions of cells as input, making it unsuited for samples with limited material - not to mention in single cells. Secondly, the method is laborious and requires many specialized protocol steps including UV cross-linking and immunoprecipitation. We here present our new method agoTRIBE, which circumvents these limitations. We show that our method yields results that are consistent with the more laborious CLIP-seq method. Importantly, the identified miRNA targets are supported by evolutionary sequence conservation and are predictors of functional miRNA repression. In addition, we show that agoTRIBE can be applied to detect miRNA-target interactions in human single cells and to deconvolute miRNA targeting in distinct phases of the cell cycle without the use of physical cell sorting.

To develop our new method, we leveraged on the TRIBE approach^11^ in which an RNA-binding protein of interest (in our case Argonaute2) is fused to the RNA-editing domain of ADAR2. The RNA-binding protein part leads the fusion protein to its natural targets, and the editing domain deaminates adenosines to inosines (A>I) in the RNA target, in effect leaving nucleotide conversions that can be detected by sequencing as A>G substitutions (Figure 1). These substitutions can in principle be detected by single-cell RNA sequencing, and the method avoids lossy isolation since it does not use immunoprecipitation. To tailor agoTRIBE for Argonaute proteins, we made three modifications to the original TRIBE approach: (1) we used a hyperactive version of the ADAR2 deaminase domain, in which a E488Q substitution results in increased editing^12, 13^; (2) we connected the Argonaute2 and ADAR2 domain with a 55 amino acid long flexible linker and (3) we fused the ADAR2 domain to the N-terminus of Argonaute2, since the protein structure of Argonaute2 indicates that fusing to the C-terminus would be detrimental to the binding to the guide miRNAs^14-18^ (Figure 1b). We confirmed that tagging Argonaute2 with the ADAR2 editing domain does not change its cytoplasmic localization (Figure 1c) or its colocalization with TNRC6B, a P-body marker (Supplementary Figure 1a). In particular, we found that agoTRIBE partly locates to cytoplasmic foci that are similar to P-bodies, which are known to be interconnected with miRNA function^19-21^ (Figure 1c, indicated by white arrows).

**Figure 1:**
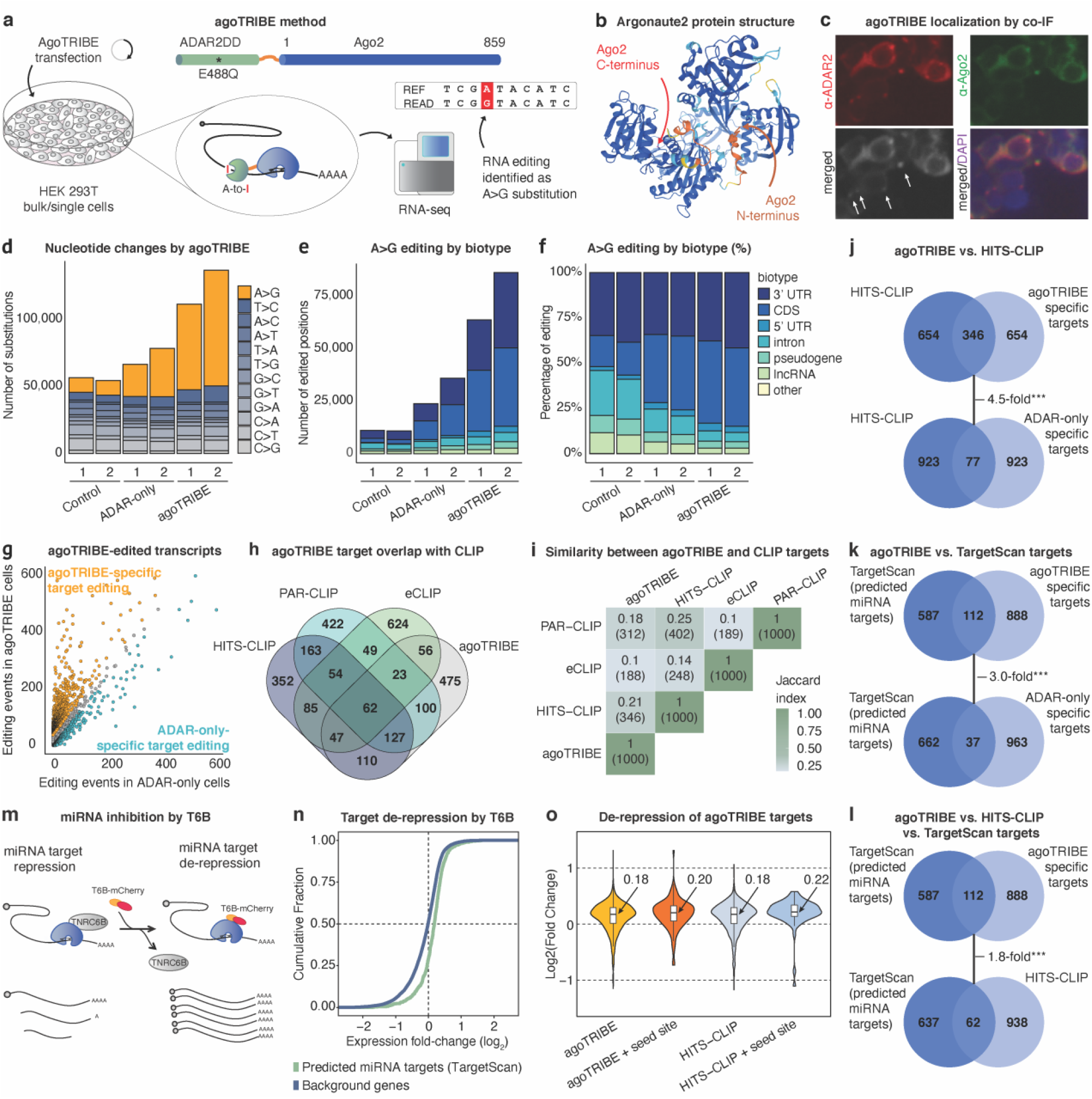
agoTRIBE detects miRNA targets through RNA editing. **a**, Schematic representation of the agoTRIBE method. The agoTRIBE approach fuses human Argonaute2 with the adenosine deaminase domain of human ADAR2, carrying a hyperactive mutation (E488Q) and depositing edits on the targeted transcripts. The edited nucleotides can be detected as A>G substitutions by standard or single-cell RNA sequencing. Above, a schematic representation of the N-terminal tagging of human Argonaute2. **b**, Human Argonaute2 protein structure prediction using AlphaFold2. Tagging of the C-terminus, which is embedded in the protein structure, could result in a misfolded protein unable to load miRNAs. **c**, Immunofluorescence staining to visualize agoTRIBE. AGO2 (green) and ADAR2 (red) co-staining was used to detect agoTRIBE while DAPI (Magenta) was used for nuclear staining. The co-localization in cytoplasmic foci (white arrows) suggests that agoTRIBE is present in P-bodies **d**, A>G editing (in orange), indicative of ADAR2 editing, specifically increases in agoTRIBE transfected cells compared to control cells or cells transfected with the ADAR2 editing domain without Argonaute2 (‘ADAR-only’ controls). **e**, Editing increases specifically in 3’ UTR and coding sequence (CDS) and remains constant in other transcript types. **f**, As in **e**, but represented as percentages of editing. **g**, Editing in agoTRIBE-transfected cells vs. ADAR-only transfected cells. Each dot corresponds to one gene. Genes in orange have substantially more editing in agoTRIBE cells while the genes in light blue have substantially more editing in ADAR-only cells (Methods). The cells in grey have comparable editing in the two conditions. **h**, Venn diagram of top 1000 miRNA targets reported by each of agoTRIBE, HITS-CLIP, PAR-CLIP and eCLIP. **i**, Jaccard similarity index values of top target sets reported by each of the four methods. **j-k**, Venn diagrams of overlaps between targets reported by agoTRIBE; targets reported by HITS-CLIP; targets specific to ADAR-only controls; and 699 miRNA targets predicted by TargetScan. **m**, Schematic representation of global miRNA inhibition by T6B. **n**, De-repression of miRNA targets predicted by TargetScan upon T6B-mCherry transfection. **o**, Increase in expression of agoTRIBE and HITS-CLIP targets upon T6B transfection. ‘Seed site’ means that the transcript harbors a conserved binding site for one of the ten most abundant miRNAs in HEK-293T cells.

Importantly, when we transiently express agoTRIBE in ∼50,000 human HEK-293T cells (Methods), we observe that A>G nucleotide substitutions – as expected by ADAR2-mediated editing - increase substantially compared to control cells (Figure 1d, Suppl. Table 1, Suppl. Table 2). In contrast, cells transfected with only the ADAR2 deaminase domain without Argonaute2 - henceforward referred as ‘ADAR-only’ - increase only moderately the number of A>G substitutions, thus suggesting the importance of miRNA guidance for the newly detected editing (Figure 1d). Of note, other types of nucleotide substitutions remain largely unchanged between the analyzed conditions, indicating specific ADAR2-mediated editing. Besides, we observe that editing in mRNA exonic regions specifically increases while editing in intronic regions and non-coding transcripts such as lncRNAs and pseudogenes remains constant (Figure 1e-f). This is consistent with miRNAs targeting mature mRNAs in the cytoplasm, while there is little evidence of miRNAs targeting non-translating sequences such as introns or long non-coding RNA transcripts^22^, which are most commonly located in the nucleus. In summary, we observe highly increased editing in mRNA transcripts that likely correspond to cytoplasmic miRNA targets.

To discern miRNA-guided editing from background editing – including that of endogenously expressed ADAR2 – we compared editing patterns upon agoTRIBE transfection relative to ‘ADAR-only’ controls (Figure 1g). We assumed that transcript editing that is specific to agoTRIBE is guided by miRNAs, while editing that is specific to ADAR-only represents background activity. To do so, we compared the total number of editing events per gene in the two conditions - agoTRIBE vs. ADAR-only - and focused on the top 1000 putative mRNA targets that showed specifically increased editing upon the agoTRIBE transfection (Figure 1g). Since Argonaute CLIP-seq represents the current state-of-the-art in experimental detection of miRNA-target interactions, we first compared our list to targets identified by different variations of the CLIP-seq methodology^23-25^. We found that the agoTRIBE top 1000 targets have substantial overlap with CLIP-seq targets, and do not represent an outlier group with an excess of unique targets not found by the CLIP-seq methods (Figure 1h). In fact, the eCLIP method reports more targets (624 unique targets) that are not shared with any other method than does agoTRIBE (485 unique targets, Figure 1h). Comparing the similarity between the target sets reported by each the four methods using the Jaccard index, we found that HITS-CLIP and PAR-CLIP resemble each other the most (Jaccard Index 0.25, Figure 1i), while agoTRIBE also has strong similarity to both of these methods (Jaccard Index 0.21 and 0.18, respectively). eCLIP in contrast had less resemblance to the other two CLIP methods (Jaccard Index 0.14 and 0.1, respectively). Importantly, more than half of the targets (525 targets) reported by agoTRIBE were supported by one or more CLIP-seq methods (Figure 1h). We considered that the overlap between methods could be due to undetected biases, for instance that highly expressed transcripts might be more efficiently detected by both CLIP-seq and agoTRIBE. We therefore compared the overlap between the top 1000 targets from agoTRIBE and HITS-CLIP with the overlap between our ADAR-only controls and HITS-CLIP (Figure 1j). Reassuringly, we found that the overlap between agoTRIBE and HITS-CLIP (346 targets) was 4.5-fold higher (p-value < 0.001, binomial test) than the overlap between our ADAR-only controls and HITS-CLIP (77 targets), indicating that the consistency between methods depends on Argonaute-guided editing and is not due to unspecific biases. Lastly, it is well-established that Argonaute CLIP can identify miRNA binding events with high resolution, in some cases to the level of the level of individual nucleotides^24^. We overlapped the editing positions of editing upon agoTRIBE transfection with reported HITS-CLIP-seq binding sites, finding little consistency in the positional information conferred by the two methods - suggesting that our method might give less precise positional information than does HITS-CLIP (Supplementary Figure 2). Overall, our results show that agoTRIBE in its reported target repertoire resembles a CLIP-seq method, even though it uses a completely distinct antibody-free approach.

**Figure 2:**
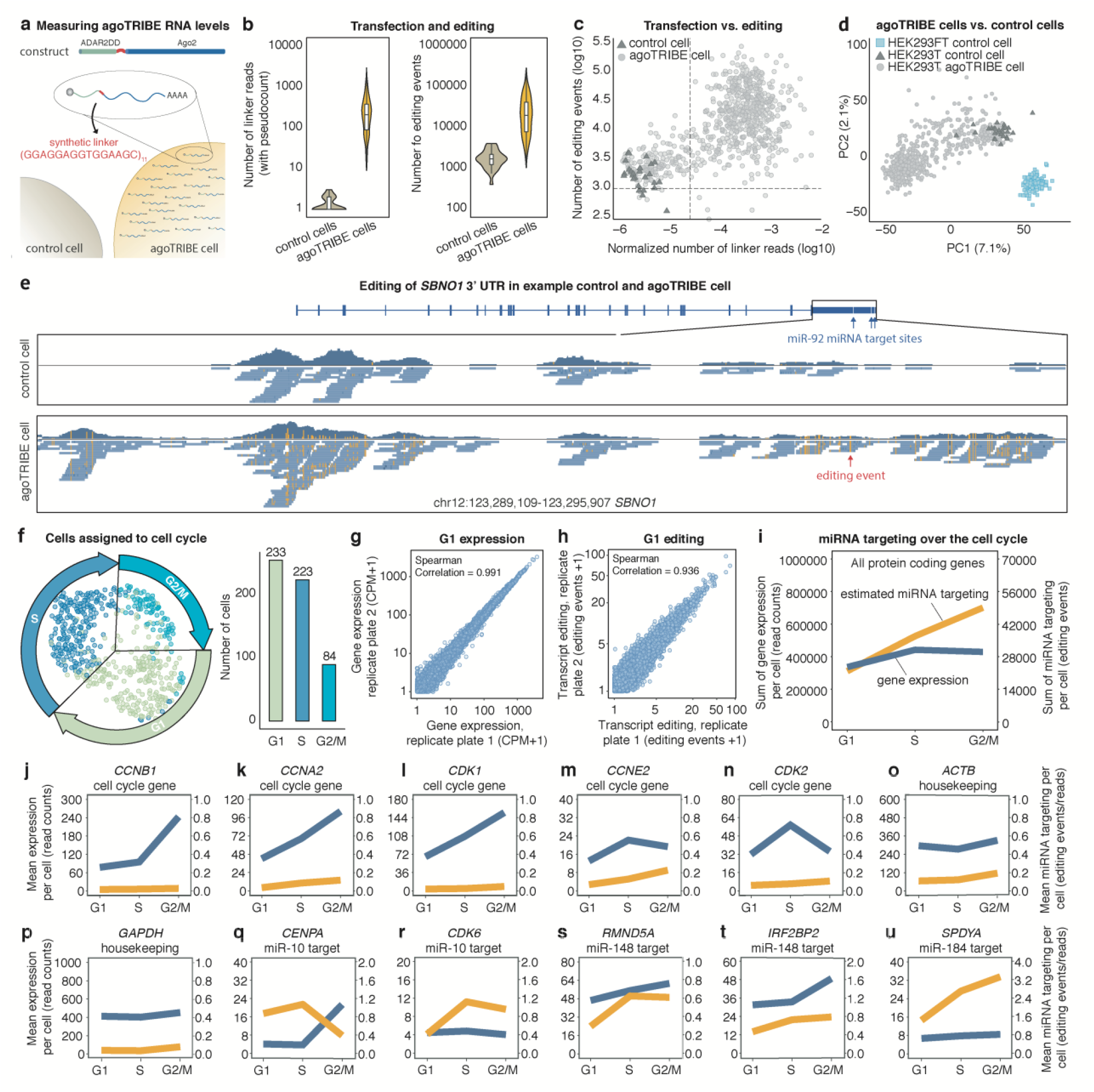
miRNA targeting in single cells. **a**, The agoTRIBE transcript includes an artificial linker that is detected in RNA sequencing and can be used to estimate agoTRIBE levels. **b**, Levels of linker sequencing reads (left) and editing events (right) detected in 703 agoTRIBE-transfected cells and 26 control cells profiled by Smart-seq3 single-cell RNA sequencing. **c**, Normalized number of linker reads vs. number of editing events for individual cells. The number of linker reads were normalized to the sequencing depth of each cell. The dotted lines indicate the thresholds for agoTRIBE cells that were used for downstream analyses. **d**, Principal component analysis of agoTRIBE transfected cells and control cells vs. control cells from a different human embryonic kidney cell line (HEK-293FT). The cells were positioned based on their Smart-seq3 transcription profiles. **e**, Editing patterns of the *SBNO1* transcript in a control cell vs. an agoTRIBE transfected cell. **f**, Overview of 540 agoTRIBE-transfected single cells assigned to cell cycle stages using their Smart-seq3 profiles. The dimensionality reduction was performed with the UMAP algorithm. **g**, Gene expression for cells assigned to the G1 cell cycle stage. The cells originate from two distinct replicate plates of single cells. Each dot indicates one gene. **h**, Transcript editing for cells assigned to the G1 cell cycle stage. **i**, Overview of estimated overall miRNA targeting during the cell cycle. Expression values and editing events were normalized to the number of cells in each cell cycle stage. **j-u**. Examples of transcript expression and estimated miRNA targeting during the cell cycle. Transcript expression is in blue and miRNA targeting in orange.

Besides experimental identification of miRNA-target interactions, computational methods to predict target sites are also available and commonly used^26-28^. These methods typically detect sequence motifs that could confer binding by specific miRNAs, and can also integrate their evolutionary conservation across species. Computational prediction inherently has high false positive and false negative rates but, on the other hand, predicted miRNA binding sites that are conserved through evolution are likely functional *bona fide* miRNA target sites. In this way, we next compared our agoTRIBE top targets to transcripts predicted to be regulated by miRNAs according to the widely used TargetScan prediction database^27^, finding an overlap of 112 transcripts (Figure 1k, Methods). This was 3-fold higher (p-value < 0.001) than the overlap with our ADAR-only control (37 transcripts) indicating that the convergence between agoTRIBE and the computational target prediction is due to Argonaute guidance. Additionally, we found that there was 1.8-fold higher overlap (p-value < 0.001) between agoTRIBE and TargetScan predictions (112 transcripts) than between HITS-CLIP and TargetScan targets (62 transcripts), providing sequence conservation evidence that agoTRIBE is more likely to report functional miRNA targets than the HITS-CLIP method (Figure 1l).

To investigate the functionality of agoTRIBE targets further, we performed a global inhibition of miRNA action in human HEK-293T cells to detect de-repressed targets. Since the TNRC6B protein is an essential cofactor for miRNA-driven post-transcriptional repression, we performed the inhibition of miRNA function by overexpressing the artificial T6B peptide, which effectively occupies the TNRC6B protein-binding pocket on Argonaute and causes global de-repression of targets^29, 30^ (Figure 1m). We indeed observed a substantial de-repression of TargetScan predicted miRNA targets, suggesting that our perturbation experiment was successful (Figure 1n, Suppl. Table 3). We next looked at the changes in gene expression of targets reported by agoTRIBE and HITS-CLIP and found that both target sets showed an 0.18 log2-fold increase in expression when miRNA function was inhibited (Figure 1o). This increased to 0.20 and 0.22 log2-fold increases for the respective methods when only targets supported by sequence motif conservation were considered (Methods). These ∼15% increases in expression are consistent with previous studies that report ∼30% de-repression at the RNA and protein levels when miRNAs are genetically deleted^31, 32^. Furthermore, miRNAs have been proposed to have subtle functions in canalizing gene expression by buffering expression noise^33, 34^. In summary, our experiment demonstrates that agoTRIBE predicts functional miRNA targets as well as does a state-of-the-art CLIP method.

The agoTRIBE approach tested here expresses an ADAR2 deaminase domain with Argonaute2 as a fusion protein from a single construct. To test the robustness of agoTRIBE, we also benchmarked two distinct designs (Supplementary Figure 2). The first relies on simultaneous expression and dimerization in living cells of an EGFP-tagged Argonaute2 and a GFP-nanobody fused with ADAR2 editing domain^35^. The second design consists of synthetic coiled-coil E3- and K3-tags, which enable to heterodimer formation when Argonaute2 and the ADAR2 editing domain are individually co-expressed in living cells^36^. We find that both approaches increase endogenous editing similarly to our fusion protein approach, demonstrating the robustness of agoTRIBE (Supplementary Figure 3). In particular, using modular designs such as the nanobody and heterodimer approach gives flexibility to experiments and allow an easy exchange of modifying domains (e.g., TRIBE^11^, hyperTRIBE^12, 13^ and STAMP^37, 38^) and RNA-binding proteins of interest.

To test the limits of the sensitivity of agoTRIBE, we transfected our construct into HEK-293T cells; sorted individual cells and subjected them to Smart-seq3 single-cell RNA sequencing^39^. In total, we profiled the transcriptomes of 703 agoTRIBE-transfected cells and 26 control cells after stringent quality controls (Methods, Suppl. Table 4, Suppl. Table 5). Since the agoTRIBE construct includes an artificial linker region, which is sequenced along with the transcriptomes, we could use sequence reads that map to the linker as an estimate of transfection efficiency of individual cells (Figure 2a). As expected, many linker reads were detected in the agoTRIBE-transfected cells, while few linker reads, likely the result of mapping artifacts, were detected in the control cells (Figure 2b, left). Overall, transcriptome-wide editing increased substantially in the agoTRIBE-transfected cells, with an average of ∼32,500 editing events in each cell compared to ∼1900 editing events in the control cells (Figure 2b, right). We also found that agoTRIBE cells with few linker reads tend to have little editing (Figure 2c), suggesting that some cells might not be efficiently transfected. These cells could however easily be computationally identified and discarded, leaving a total of 540 efficiently transfected and edited cells for downstream analyses (Figure 2c, dotted lines). Importantly, we found that the transcriptional profiles of the agoTRIBE-transfected cells overall resemble those of control cells (Figure 2d, light and dark grey), suggesting that editing by our fusion protein does not substantially alter the transcriptome composition - even in measurements with sensitive single-cell methods such as the Smart-seq3 protocol. In contrast, control cells belonging to the fast-growing HEK-293FT cell line, but sequenced with the same protocol, clearly cluster separately from our control and agoTRIBE cells (Figure 2d, in light blue).

We observed that some specific miRNA targets increase strongly in editing. For instance, *SBNO1* has three binding sites for miR-92, which is a highly expressed miRNA in HEK-293T cells (Figure 2e, blue arrows). The density plots of sequenced transcript parts show that the sensitive Smart-seq3 protocol yields transcript information for much of the 3’ UTR in a control cell and an agoTRIBE-transfected cell (Figure 2e, blue densities). Editing is virtually absent in the control cell but prevalent in the agoTRIBE cell (Figure 2e, editing in orange). As a proof-of-principle of biological applications of our methods, we next applied Seurat^40^ to computationally sort the 540 agoTRIBE-transfected single cells into the G1, S and G2/M stages of the cell cycle (Figure 2f). Even when stratifying the cells into distinct cell cycle stages, our measurements of gene expression (Figure 2g) and editing (Figure 2h) were still highly reproducible. We found that overall normalized gene expression stays constant over the cell cycle, while miRNA targeting events, as measured by global editing patterns, increases throughout the cell cycle from G1 to M (Figure 2i). This is consistent with previous observations that miRNA repression is weakest in the G1 phase^41^, but the increase in editing could also represent accumulated miRNA targeting over the cell cycle, which is then diluted by new transcription in the G1 phase^42, 43^. We find that known cell cycle-specific genes behave as expected in our single-cell data (Figure 2j-n). For instance, *CCNB1* has the highest expression in the G2/M stage (blue line), consistent with its role in promoting the transition from G2 to the mitosis phase of the cell cycle (Figure 2j). Similarly, *CDK2* is predominantly expressed in the S stage, consistent with its role in progressing through the G1-S checkpoint (Figure 2n). These specific cell-cycle genes do not appear to be regulated by miRNAs, as evidenced by their low levels of editing across the entire cell cycle (yellow lines). In contrast, we found numerous genes that are differentially targeted by miRNAs across the cell cycle. For instance, the transcript of the centromeric protein CENPA appears to be strongly targeted by miRNAs during the G1 and S stages, but this targeting seems alleviated in G2/M, where the expression increases strongly, consistent with its role in mitosis (Figure 2q). The *CDK6* gene has important roles in the G1-S transition, and, interestingly, we find it to be specifically targeted in the S stage, where its expression might not be required (Figure 2r). These examples serve as a proof-of-principle that agoTRIBE can detect miRNA targeting both in single cells and across distinct populations of single cells.

In summary, we here present the first method to detect miRNA-target interactions transcriptome-wide in single cells. In a comparison with the current state-of-the-art Argonaute CLIP-seq methods, we find that agoTRIBE has several advantages (Table 1). Firstly, agoTRIBE does not require the use of antibodies, but rather simple transfection – thus reducing the cost and time of the required experimental procedures by several days. Secondly, our method uses either ordinary bulk RNA-seq or single-cell RNA sequencing to detect editing events, meaning that transcriptome-wide measurements of RNA levels are also provided as part of the protocol. Indeed, agoTRIBE transfection is so straight-forward that it could be applied to any given standard RNA-seq experiment to provide transcriptome-wide miRNA-target interaction information at little additional cost or effort. Thirdly, while Argonaute CLIP-seq requires millions of cells, agoTRIBE can be applied to individual cells. This will allow us to study heterogeneity of miRNA targeting in homogenous cell populations, about which little is currently known. It will also allow us to study miRNA targeting in complex cell compositions in cell culture, for instance in organoids or during induced cell differentiation. Importantly, the agoTRIBE fusion protein could be placed under an inducible promoter in a living organism, which would allow the profiling of miRNA targeting in individual cell types of a complex tissue, such as the mouse brain. Since the editing events would be detected with single-cell RNA sequencing, the individual cells could be sorted into cell types using computational approaches, and would not require any physical sorting. In conclusion, we foresee that agoTRIBE has numerous applications that will benefit the wider community and that it will facilitate the entry of the miRNA field into the single-cell era.

**Table 1:** Comparison between Argonaute CLIP-seq methods and agoTRIBE.

| Method | Argonaute CLIP-seq | agoTRIBE |
| --- | --- | --- |
| Approach | Immunoprecipitation in lysed cells | Editing in living cells |
| Antibodies | Needed | Not needed |
| Starting input | Millions of cells | Down to single cells |
| Specialized lab equipment | Needed | Not needed |
| Duration of protocol (excluding cell work) | >4 days | 1.5-2 days |
| Exact miRNA binding positions | Available | Not available |
| Extra information | - | Provides RNA-seq data |

## Acknowledgements

Smart-seq3 was performed at the SciLifeLab Eukaryotic Single-Cell Genomics (ESCG) facility. Cell sorting was performed at the Biomedicum flow cytometry core facility (BFC), the Karolinska Institute. Computation was performed at the Swedish National Infrastructure for Computing (SNIC) at UPPMAX. We thank members of the Friedländer, Kutter and Pelechano labs for insightful suggestions. We especially thank Dr. Albin Widmark for his feedback on human ADAR2 editing activity and sharing the ADAR2 constructs (a gift from Mary O’Connell, Masaryk University, Brno, Czech Republic), and Dr. Alexey Amunts for his helpful feedback on 3D-protein structure prediction by AlphaFold2 and PyMOL.

## Contributions

M.R.F and I.B. conceived the study. I.B. designed and developed the agoTRIBE fusion and modular versions; performed protein structure prediction and all lab experiments except where specified below. V.S. performed all computational analyses, except where specified below. E.M-S. and P.K. performed the T6B-mCherry analyses. P.K. in addition made substantial contributions to the single-cell analyses and E.M-S. supervised on editing pipeline development and gene annotation curation. L.S. advised on the nanobody design and experiments. E.A.S. advised on RNA editing analyses. E.D. assisted with benchmarking of editing analyses and F.B. designed and cloned the pilot version of the fusion protein. M.R.F. supervised all computational analyses. M.R.F. and I.B. wrote the manuscript with input from all authors.

## References

1. Bartel, D.P. Metazoan MicroRNAs. Cell 173, 20–51 (2018).

2. Valencia-Sanchez, M.A., Liu, J., Hannon, G.J. & Parker, R. Control of translation and mRNA degradation by miRNAs and siRNAs. Genes & development 20, 515–524 (2006).

3. Chekulaeva, M. & Filipowicz, W. Mechanisms of miRNA-mediated post-transcriptional regulation in animal cells. Curr Opin Cell Biol 21, 452–460 (2009).

4. Jonas, S. & Izaurralde, E. Towards a molecular understanding of microRNA-mediated gene silencing. Nature reviews. Genetics 16, 421–433 (2015).

5. Gebert, L.F.R. & MacRae, I.J. Regulation of microRNA function in animals. Nature reviews. Molecular cell biology 20, 21–37 (2019).

6. Bartel, D.P. & Chen, C.Z. Micromanagers of gene expression: the potentially widespread influence of metazoan microRNAs. Nature reviews. Genetics 5, 396–400 (2004).

7. Flynt, A.S. & Lai, E.C. Biological principles of microRNA-mediated regulation: shared themes amid diversity. Nature reviews. Genetics 9, 831–842 (2008).

8. Sun, K. & Lai, E.C. Adult-specific functions of animal microRNAs. Nature reviews. Genetics 14, 535–548 (2013).

9. Hausser, J. & Zavolan, M. Identification and consequences of miRNA-target interactions--beyond repression of gene expression. Nature reviews. Genetics 15, 599–612 (2014).

10. Markus Hafner, M.K., Tino Köster, James Marks, Joyita Mukherjee, Dorothee Staiger, Jernej Ule, Mihaela Zavolan CLIP and complementary methods. Nature Reviews Methods Primers 1 (2021).

11. McMahon, A.C. et al. TRIBE: Hijacking an RNA-Editing Enzyme to Identify Cell-Specific Targets of RNA-Binding Proteins. Cell 165, 742–753 (2016).

12. Xu, W., Rahman, R. & Rosbash, M. Mechanistic implications of enhanced editing by a HyperTRIBE RNA-binding protein. Rna 24, 173–182 (2018).

13. Rahman, R., Xu, W., Jin, H. & Rosbash, M. Identification of RNA-binding protein targets with HyperTRIBE. Nat Protoc 13, 1829–1849 (2018).

14. Elkayam, E. et al. The structure of human argonaute-2 in complex with miR-20a. Cell 150, 100–110 (2012).

15. Schirle, N.T. & MacRae, I.J. The crystal structure of human Argonaute2. Science 336, 1037–1040 (2012).

16. Baek, M. et al. Accurate prediction of protein structures and interactions using a three-track neural network. Science 373, 871–876 (2021).

17. Jumper, J. et al. Highly accurate protein structure prediction with AlphaFold. Nature 596, 583–589 (2021).

18. Nakanishi, K., Weinberg, D.E., Bartel, D.P. & Patel, D.J. Structure of yeast Argonaute with guide RNA. Nature 486, 368–374 (2012).

19. Liu, J., Valencia-Sanchez, M.A., Hannon, G.J. & Parker, R. MicroRNA-dependent localization of targeted mRNAs to mammalian P-bodies. Nat Cell Biol 7, 719–723 (2005).

20. Pillai, R.S. et al. Inhibition of translational initiation by Let-7 MicroRNA in human cells. Science 309, 1573–1576 (2005).

21. Pauley, K.M. et al. Formation of GW bodies is a consequence of microRNA genesis. EMBO Rep 7, 904–910 (2006).

22. Biasini, A. et al. Translation is required for miRNA-dependent decay of endogenous transcripts. EMBO J 40, e104569 (2021).

23. Hafner, M. et al. Transcriptome-wide identification of RNA-binding protein and microRNA target sites by PAR-CLIP. Cell 141, 129–141 (2010).

24. Kishore, S. et al. A quantitative analysis of CLIP methods for identifying binding sites of RNA-binding proteins. Nature methods 8, 559–564 (2011).

25. Patel, R.K., West, J.D., Jiang, Y., Fogarty, E.A. & Grimson, A. Robust partitioning of microRNA targets from downstream regulatory changes. Nucleic acids research 48, 9724–9746 (2020).

26. Bartel, D.P. MicroRNAs: target recognition and regulatory functions. Cell 136, 215–233 (2009).

27. Agarwal, V., Bell, G.W., Nam, J.W. & Bartel, D.P. Predicting effective microRNA target sites in mammalian mRNAs. Elife 4 (2015).

28. Riolo, G., Cantara, S., Marzocchi, C. & Ricci, C. miRNA Targets: From Prediction Tools to Experimental Validation. Methods Protoc 4 (2020).

29. Hauptmann, J. et al. Biochemical isolation of Argonaute protein complexes by Ago-APP. Proceedings of the National Academy of Sciences of the United States of America 112, 11841–11845 (2015).

30. La Rocca, G. et al. Inducible and reversible inhibition of miRNA-mediated gene repression in vivo. Elife 10 (2021).

31. Baek, D. et al. The impact of microRNAs on protein output. Nature 455, 64–71 (2008).

32. Selbach, M. et al. Widespread changes in protein synthesis induced by microRNAs. Nature 455, 58–63 (2008).

33. Hornstein, E. & Shomron, N. Canalization of development by microRNAs. Nature genetics 38 Suppl, S20–24 (2006).

34. Schmiedel, J.M. et al. Gene expression. MicroRNA control of protein expression noise. Science 348, 128–132 (2015).

35. Prole, D.L. & Taylor, C.W. A genetically encoded toolkit of functionalized nanobodies against fluorescent proteins for visualizing and manipulating intracellular signalling. BMC Biol 17, 41 (2019).

36. Fernandez-Rodriguez, J. & Marlovits, T.C. Induced heterodimerization and purification of two target proteins by a synthetic coiled-coil tag. Protein Sci 21, 511–519 (2012).

37. Meyer, K.D. DART-seq: an antibody-free method for global m(6)A detection. Nature methods 16, 1275–1280 (2019).

38. Brannan, K.W. et al. Robust single-cell discovery of RNA targets of RNA-binding proteins and ribosomes. Nature methods 18, 507–519 (2021).

39. Hagemann-Jensen, M. et al. Single-cell RNA counting at allele and isoform resolution using Smart-seq3. Nature biotechnology 38, 708–714 (2020).

40. Hao, Y. et al. Integrated analysis of multimodal single-cell data. Cell 184, 3573–3587 e3529 (2021).

41. Vasudevan, S., Tong, Y. & Steitz, J.A. Cell-cycle control of microRNA-mediated translation regulation. Cell cycle 7, 1545–1549 (2008).

42. Rodriques, S.G. et al. RNA timestamps identify the age of single molecules in RNA sequencing. Nature biotechnology 39, 320–325 (2021).

43. Hsiung, C.C. et al. A hyperactive transcriptional state marks genome reactivation at the mitosis-G1 transition. Genes & development 30, 1423–1439 (2016).

